## Supplementary information for "The unusual structural properties and potential biological relevance of switchback DNA"

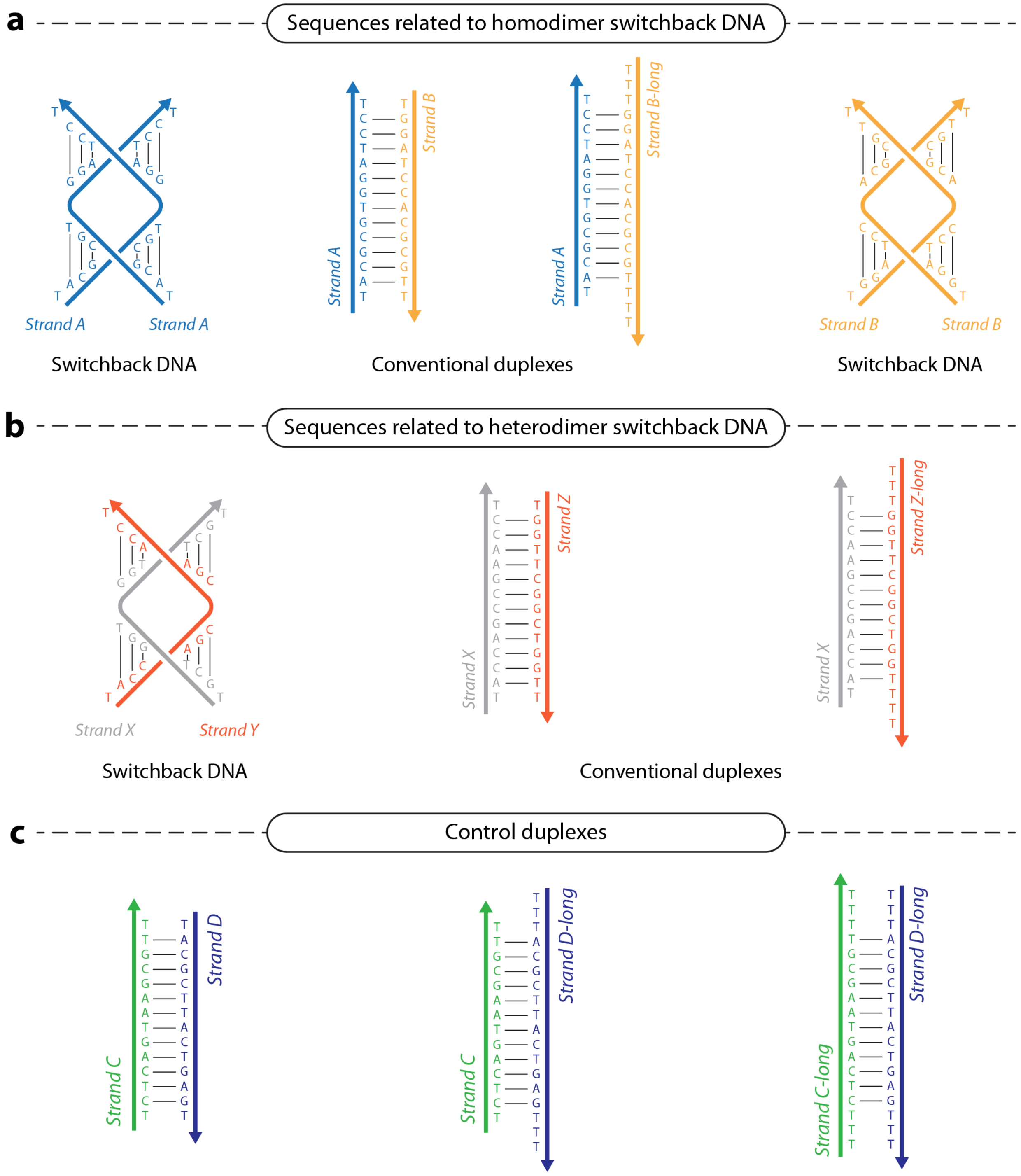


**Supplementary Fig. 1**. Sequences of DNA complexes used in this study. (a) Homodimeric switchback DNA and its corresponding conventional duplexes. (b) Heterodimeric switchback DNA and its corresponding conventional duplexes. (c) Control structures.

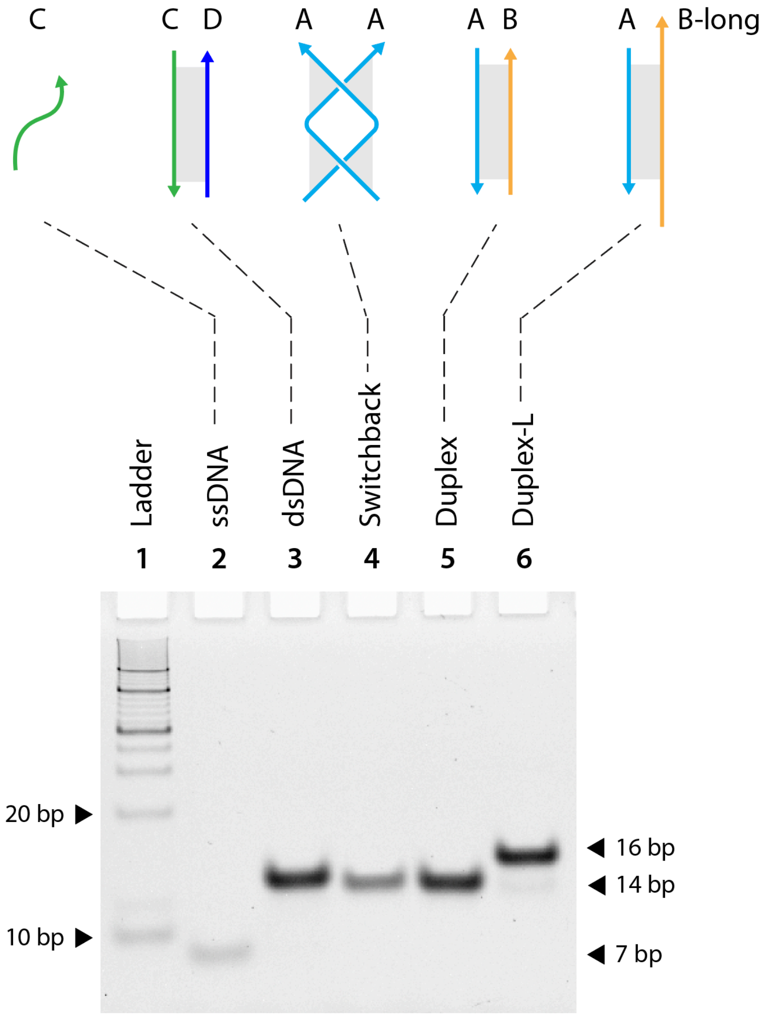


**Supplementary Fig. 2**. Non-denaturing gel showing the assembly of homodimer switchback and its corresponding duplex. Full image of the gel shown in Fig. 2a.

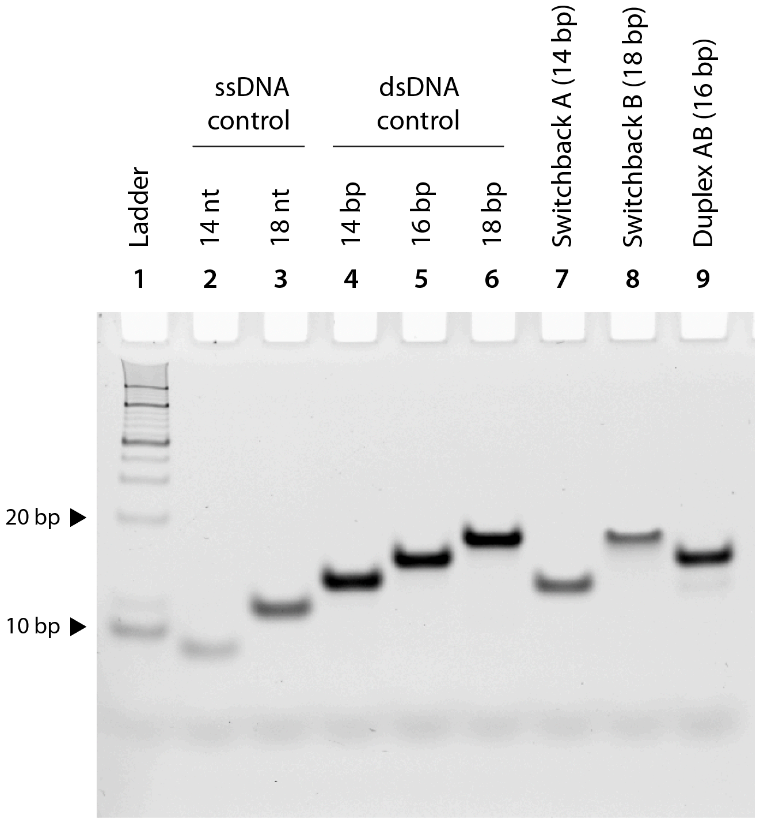


**Supplementary Fig. 3**. Non-denaturing gel showing the assembly of homodimer switchback, corresponding duplex, and control structures.

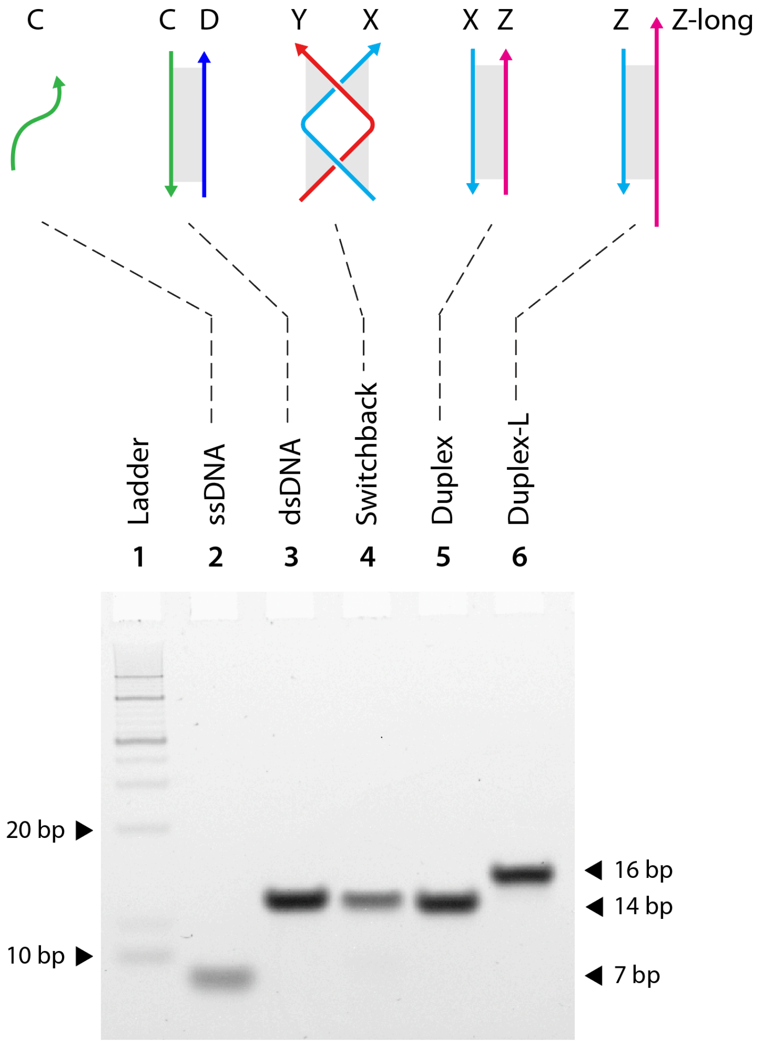


**Supplementary Fig. 4**. Non-denaturing gel showing the assembly of heterodimer switchback and its corresponding duplex. Full image of the gel shown in Fig. 2b.

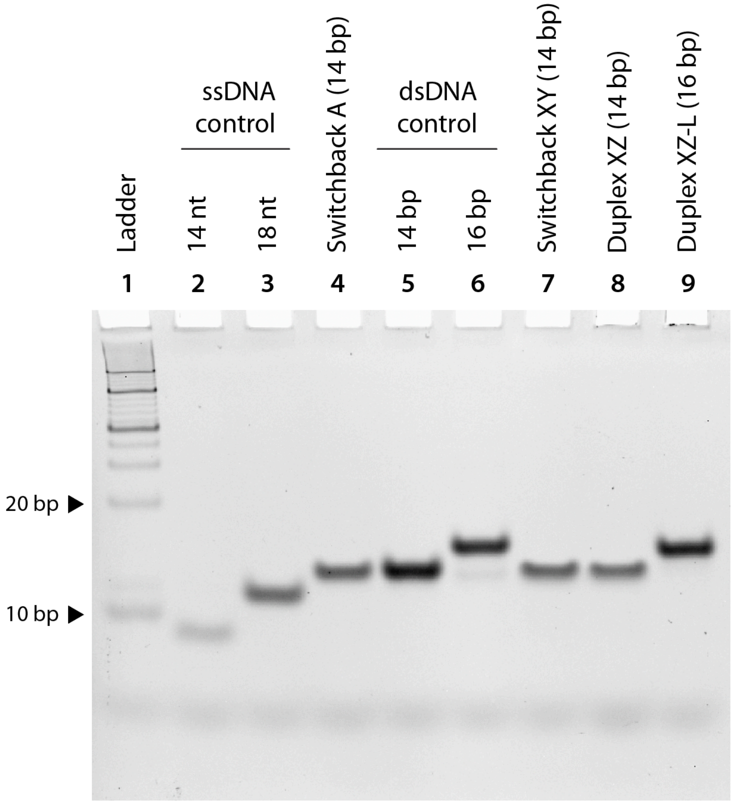


**Supplementary Fig. 5**. Non-denaturing gel showing the assembly of heterodimer switchback, corresponding duplex, and control structures.

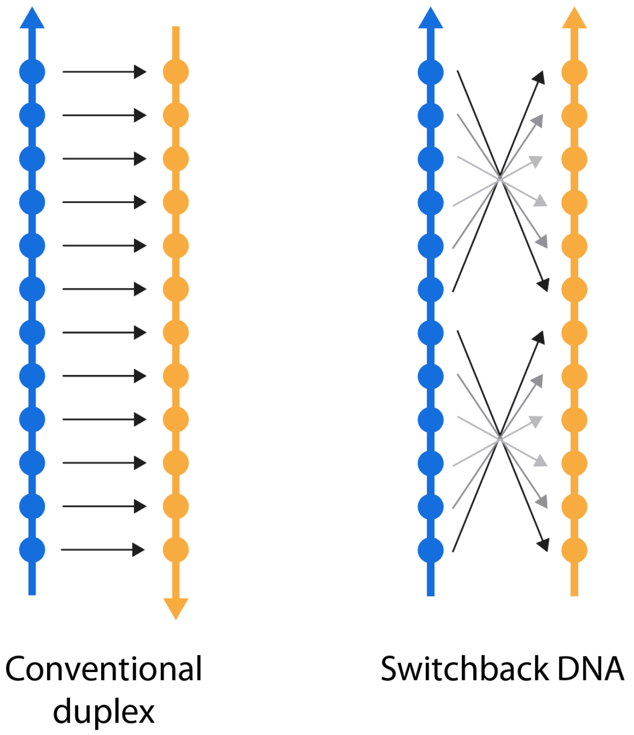


**Supplementary Fig. 6**. Base pairing rules for conventional duplexes and switchback DNA. Shown here is a structure with two half-turn domains. The black arrows map the pairs of complementary bases.

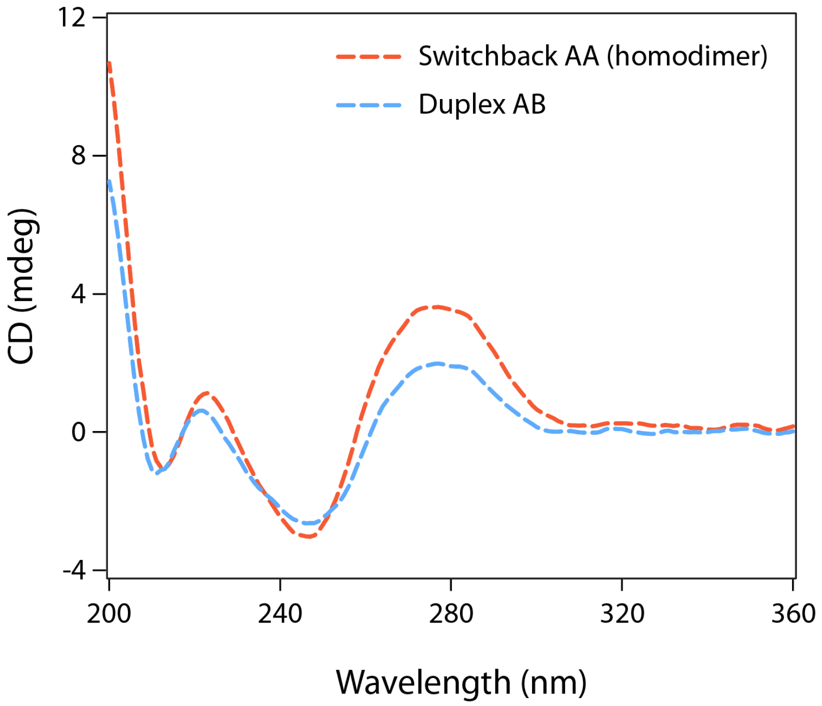


**Supplementary Fig. 7**. CD spectra of homodimeric switchback DNA and its corresponding conventional duplex.

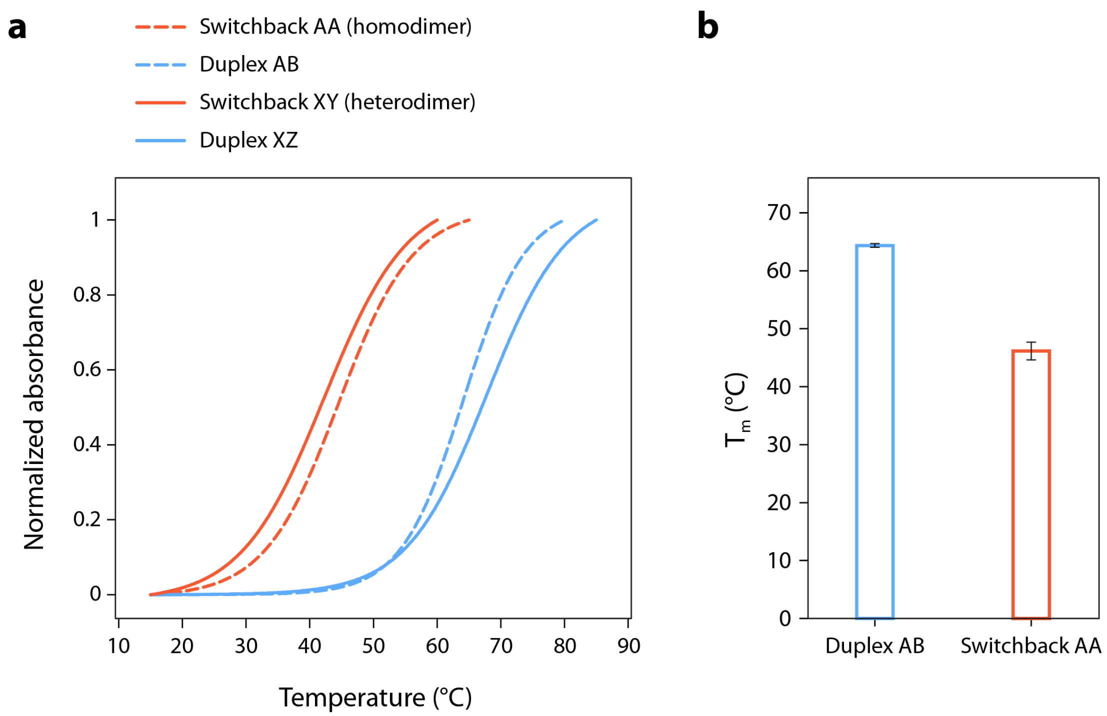


**Supplementary Fig. 8**. (a) UV melting curves of switchback DNA and conventional duplexes. (b) Melting temperatures (Tm) of homodimer switchback DNA and the corresponding conventional duplex.

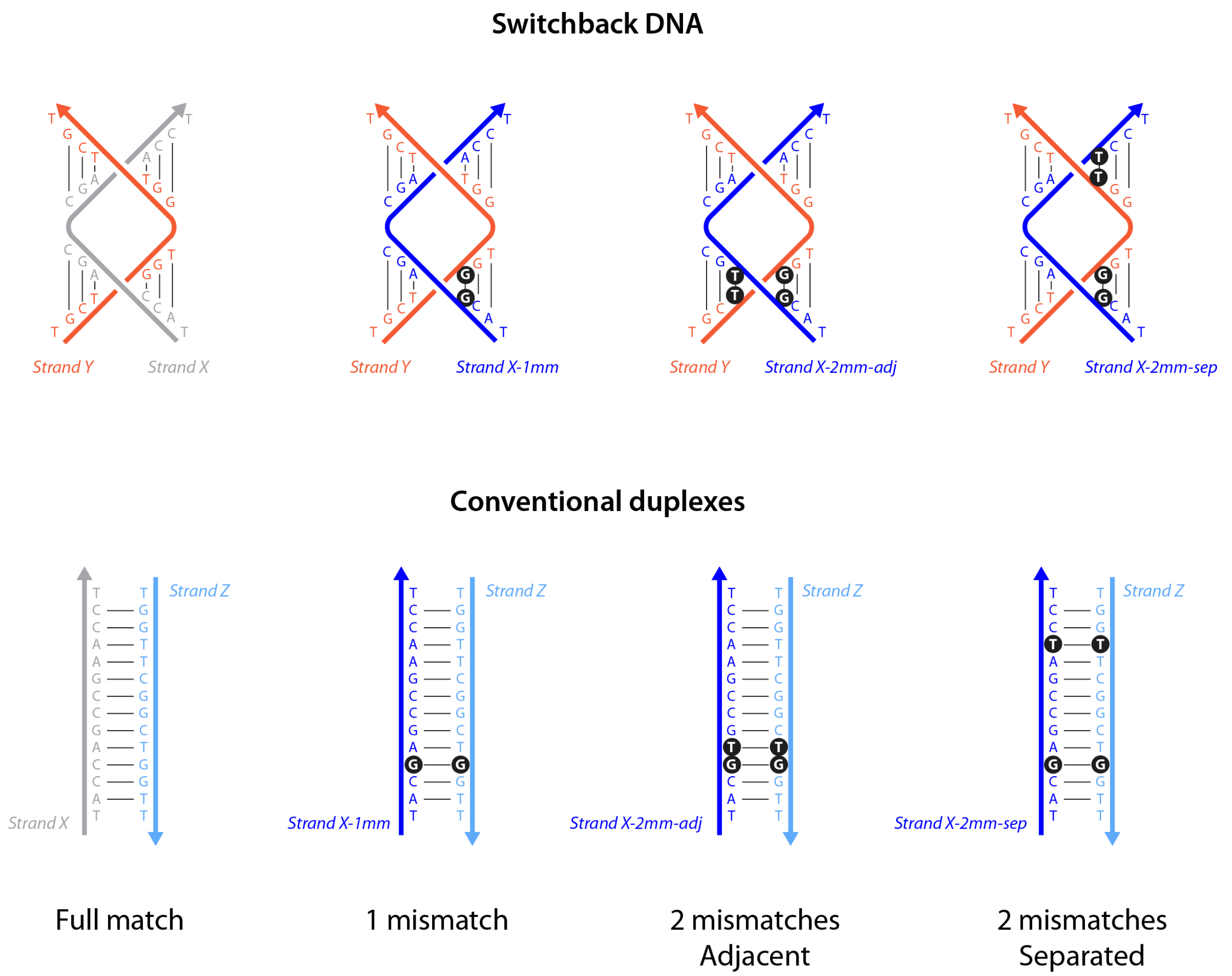


**Supplementary Fig. 9**. Design of switchback DNA and conventional duplexes with mismatched base pairs.

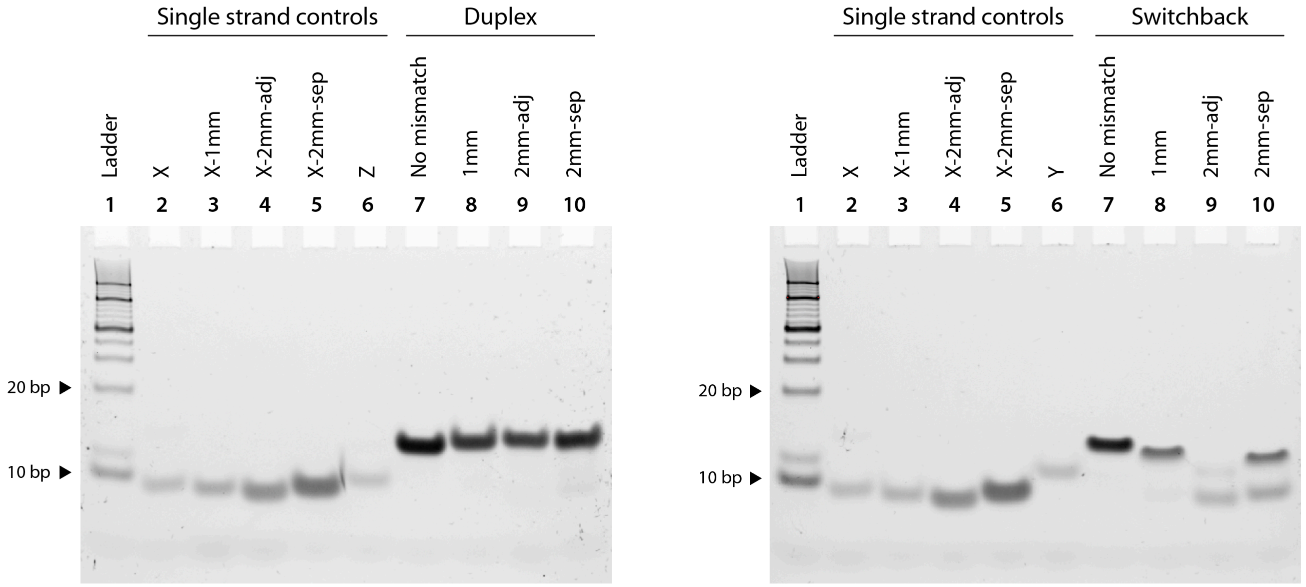


**Supplementary Fig. 10**. Non-denaturing polyacrylamide gel image of conventional duplex and switchback DNA with mismatched base pairs.

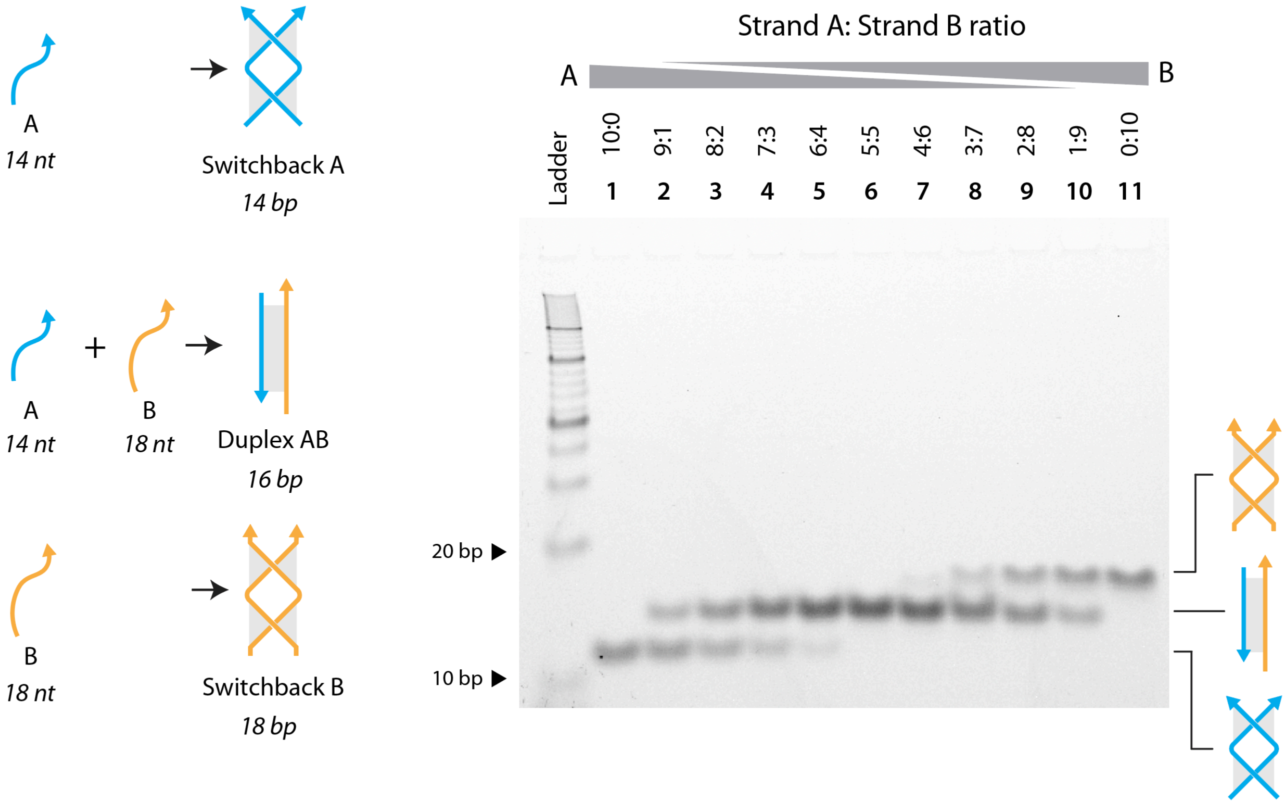


**Supplementary Fig. 11**. Strand competition between switchback DNA and conventional duplex in homodimer assembly. Full image of the gel shown in Fig. 3a.

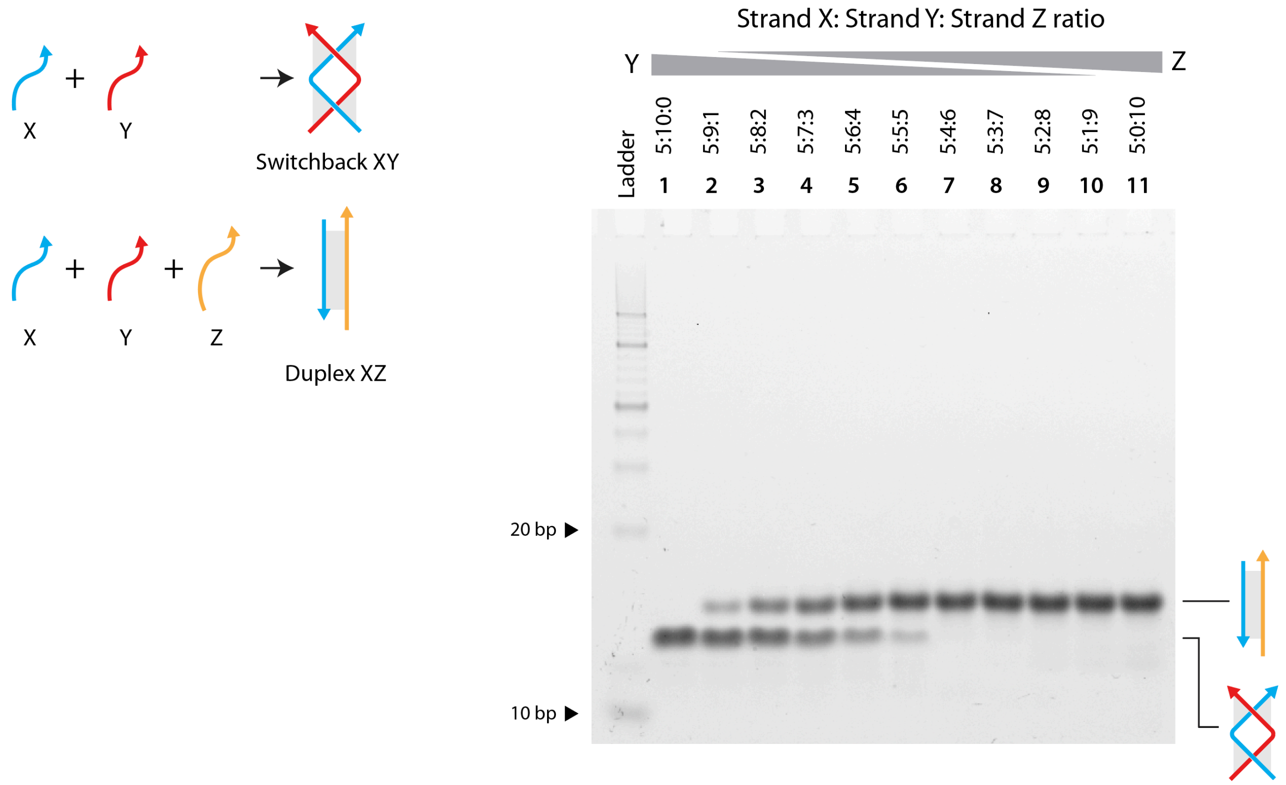


**Supplementary Fig. 12**. Strand competition between switchback DNA and conventional duplex in heterodimer assembly. Full image of the gel shown in Fig. 3b.

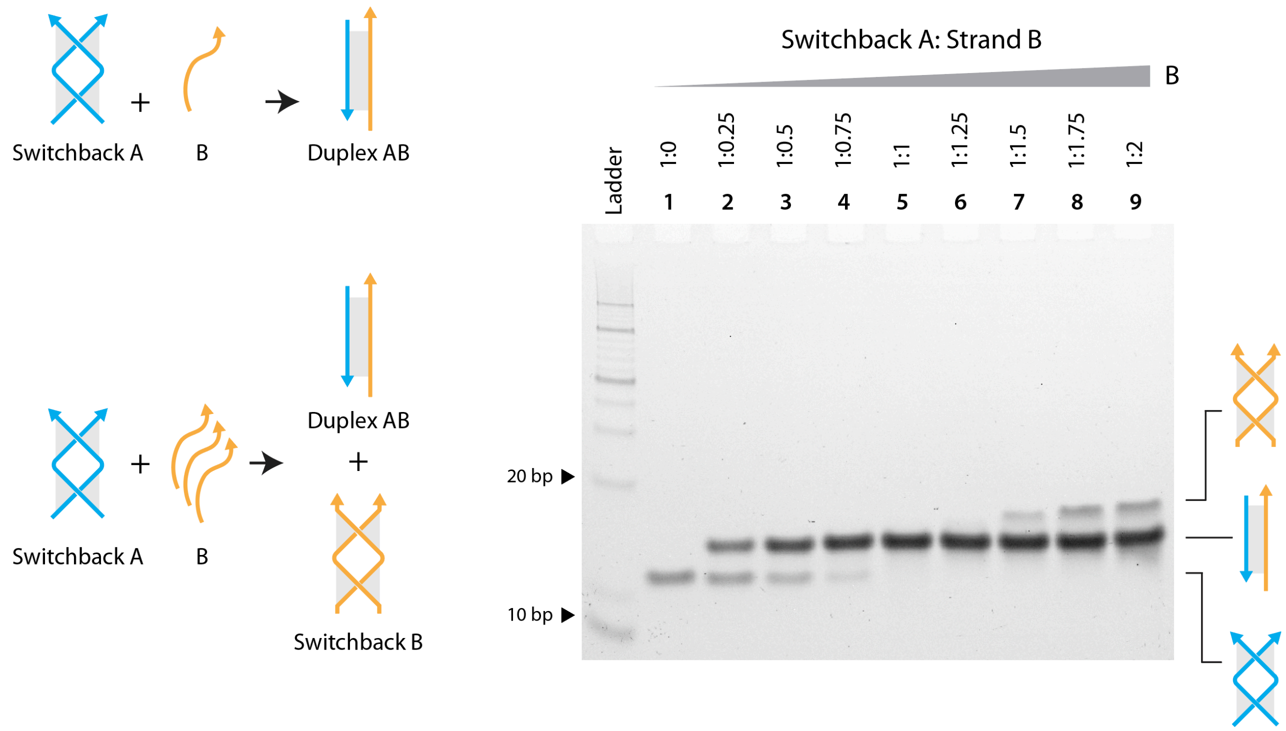


**Supplementary Fig. 13**. Strand displacement in homodimer switchback DNA on the addition of a duplex complement. Full image of gel shown in Fig. 3c.

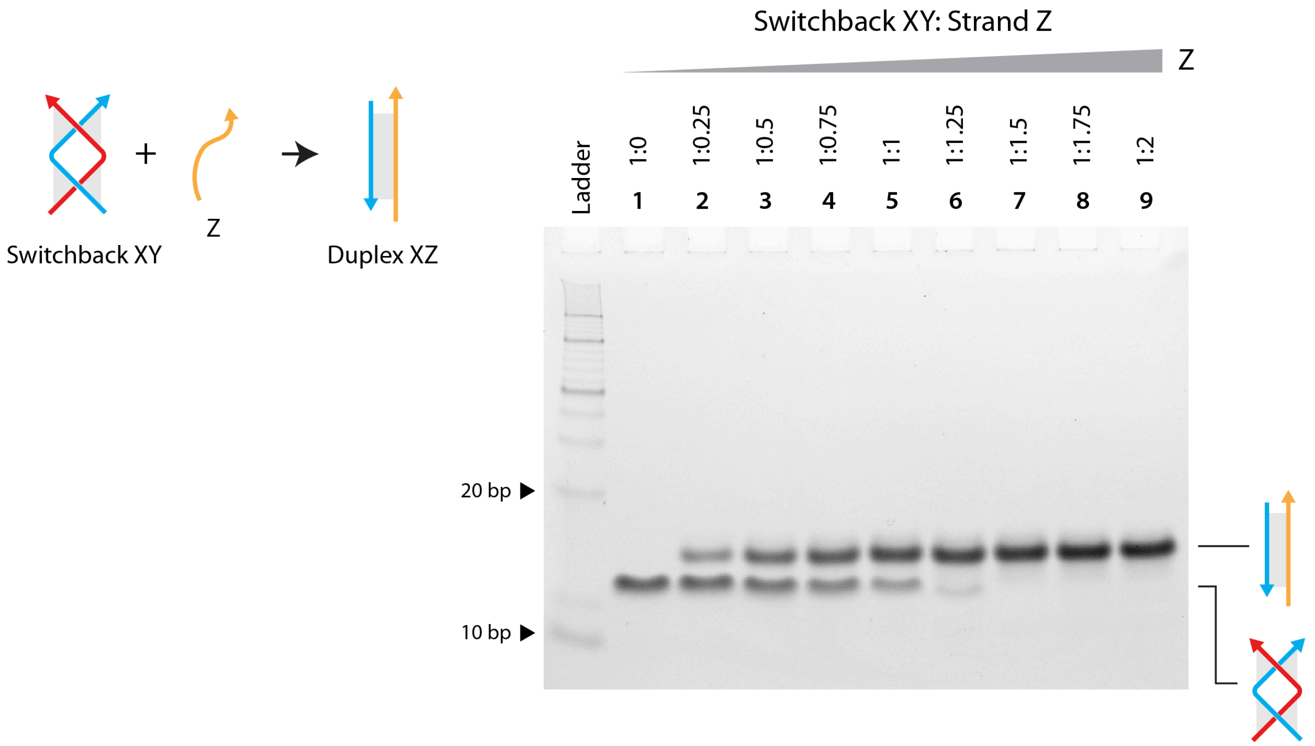


**Supplementary Fig. 14**. Strand displacement in heterodimer switchback DNA on the addition of a duplex complement. Full image of gel shown in Fig. 3d.

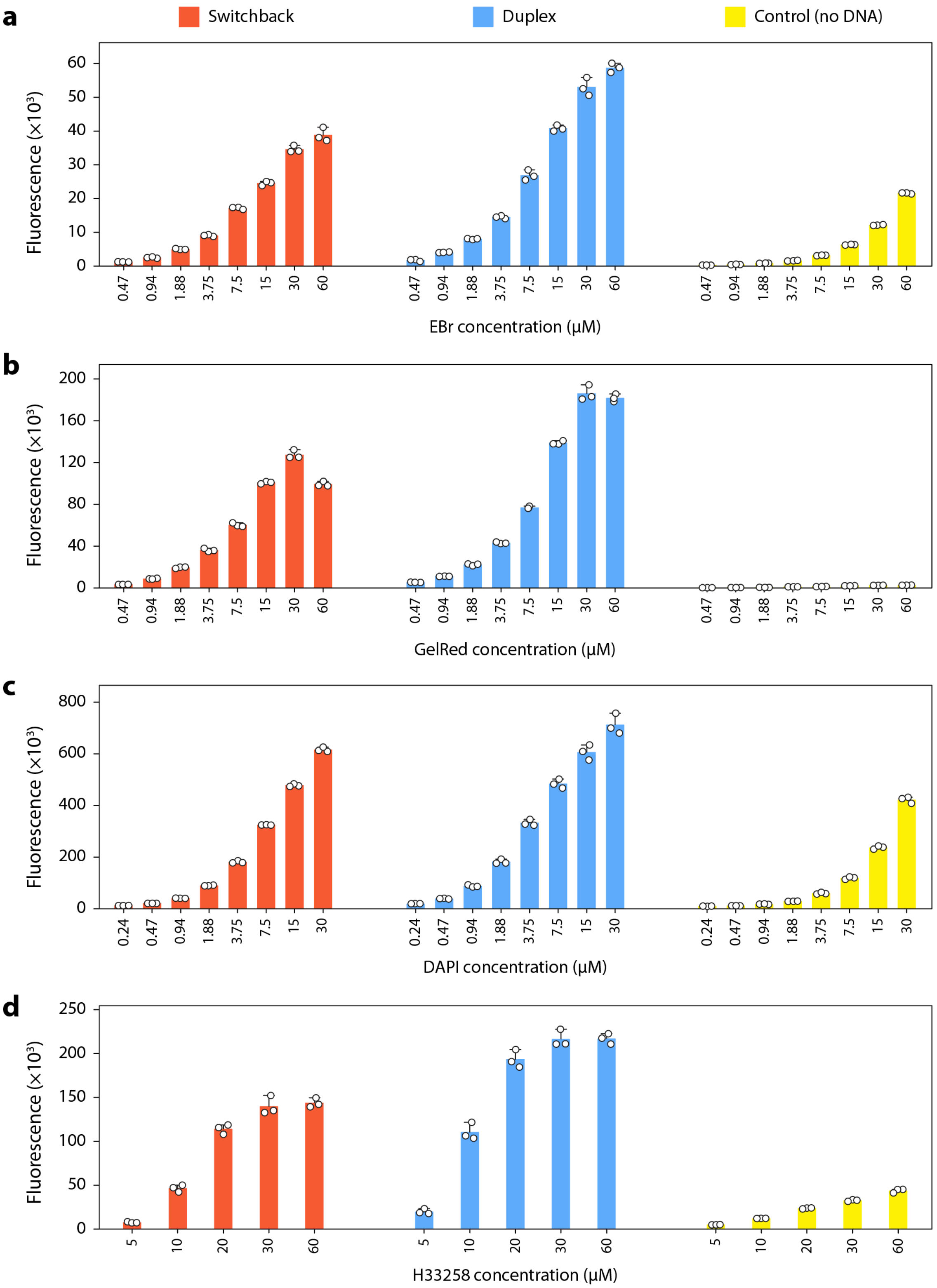


**Supplementary Fig. 15**. Raw fluorescence signals of small molecules with and without DNA.

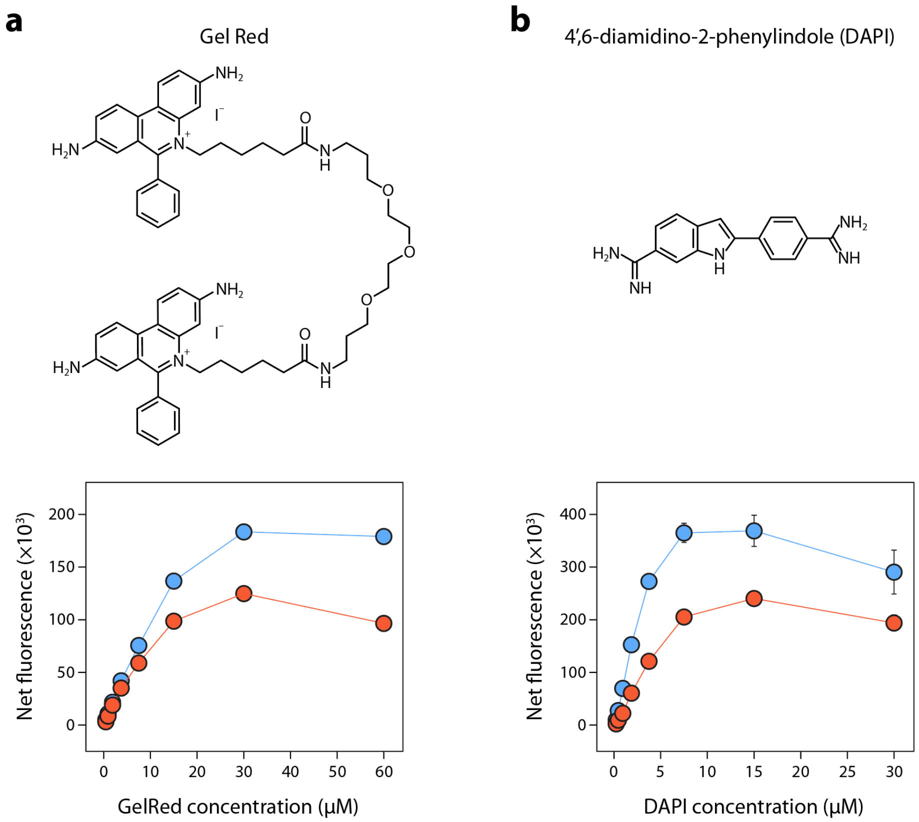


**Supplementary Fig. 16.** Fluorescence intensities of switchback DNA and conventional duplexes with different concentrations of (a) GelRed (intercalator) and (b) DAPI (minor groove binder). Data represent mean and error propagated from standard deviations of experiments performed in triplicates.

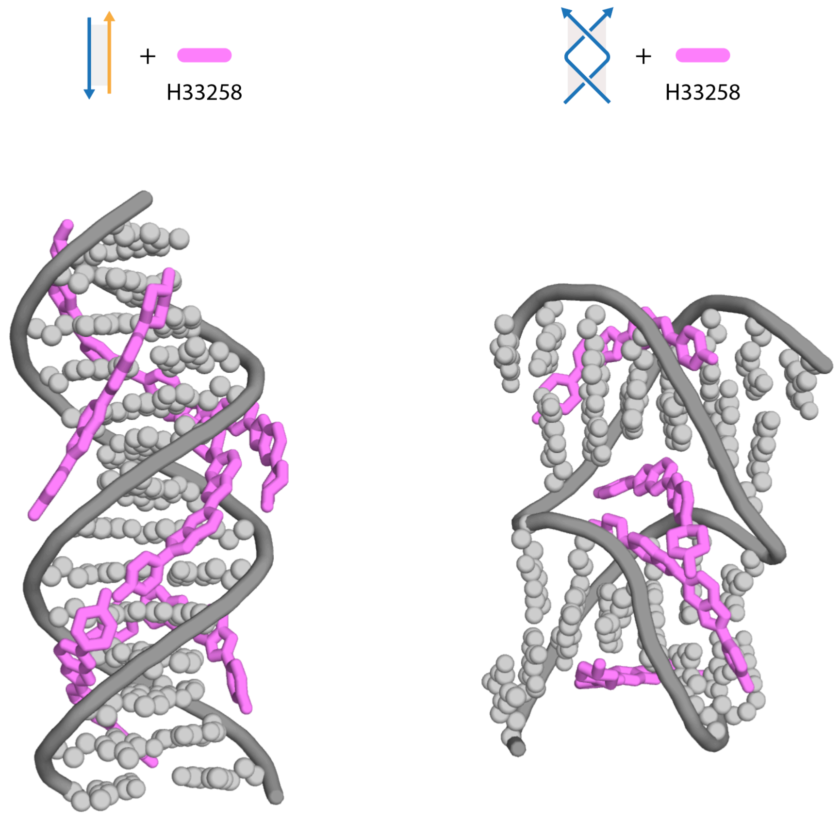


**Supplementary Fig. 17**. Molecular docking analysis of conventional duplex and switchback DNA with H33258 respectively. We sequentially docked H33258 on the duplex or switchback DNA until the number of predicted contacts between the ligand and DNA in a docking run was less than two.

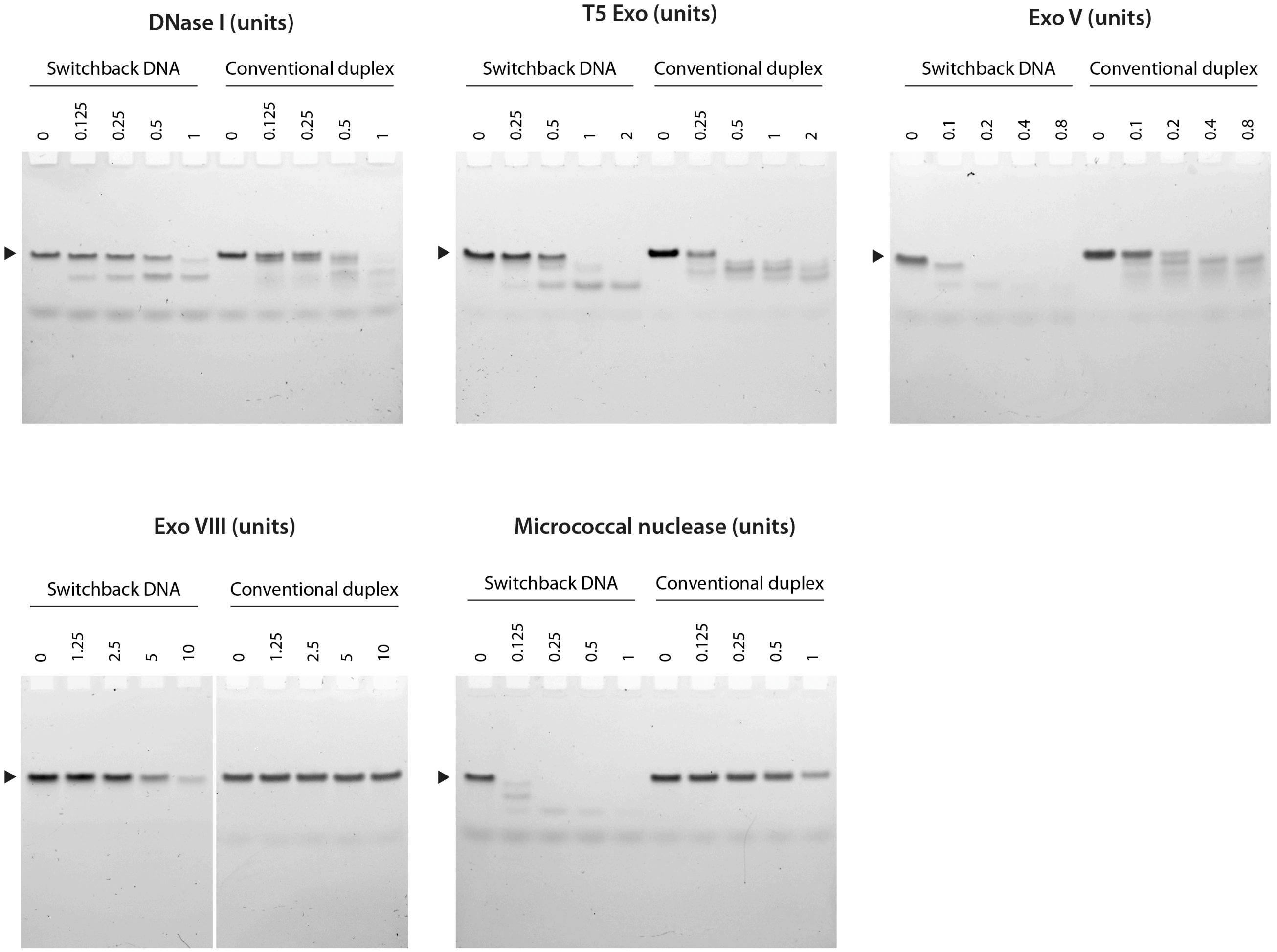


**Supplementary Fig. 18**. Non-denaturing gel images of switchback DNA and conventional duplex treated with different amounts of various nucleases.

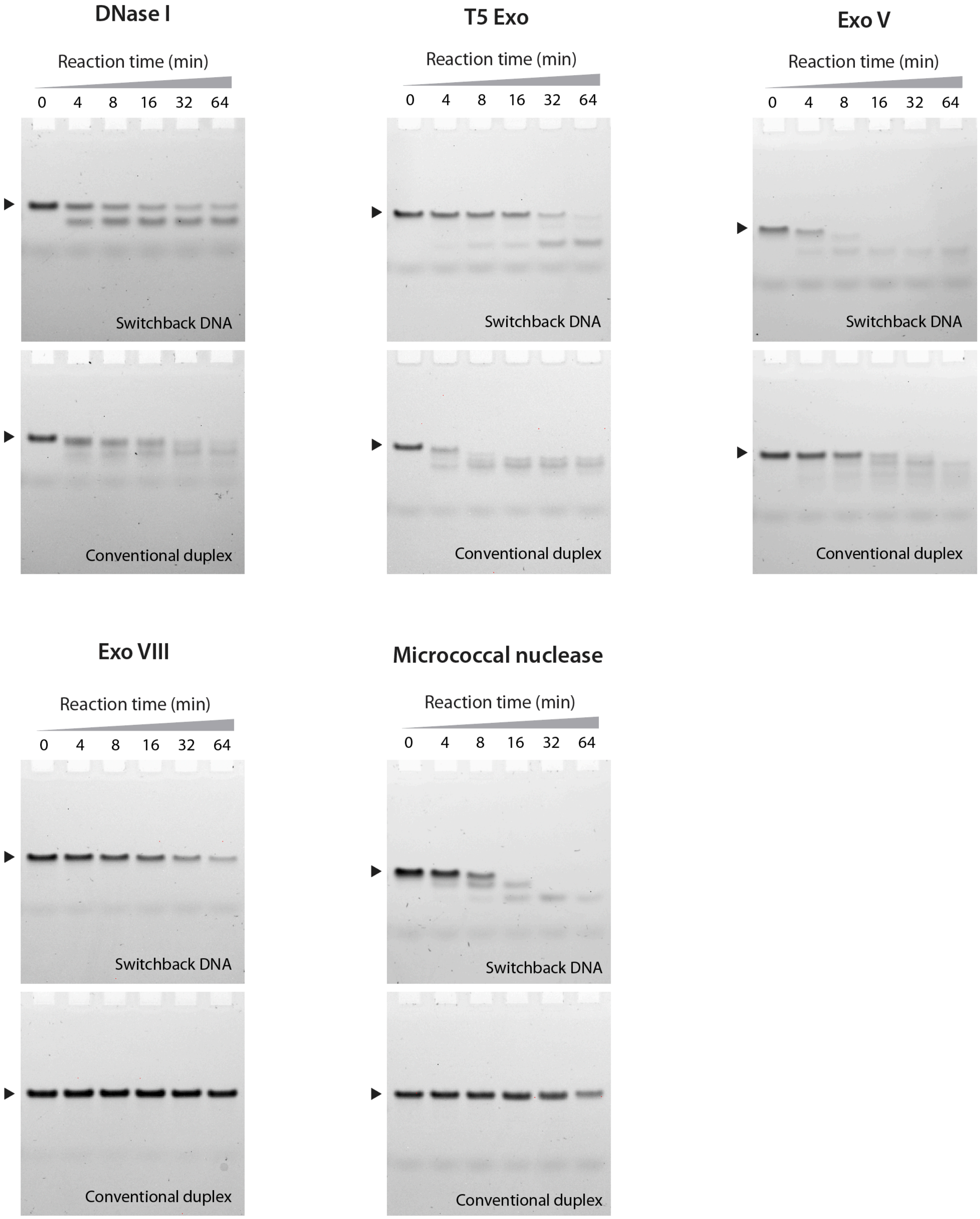


**Supplementary Fig. 19**. Non-denaturing gel images of switchback DNA and conventional duplex treated with various nucleases for different time periods (DNase I: 0.75 U, T5 Exo: 1 U, Exo V: 0.5 U, Exo VIII: 10 U, micrococcal nuclease: 0.25 U).

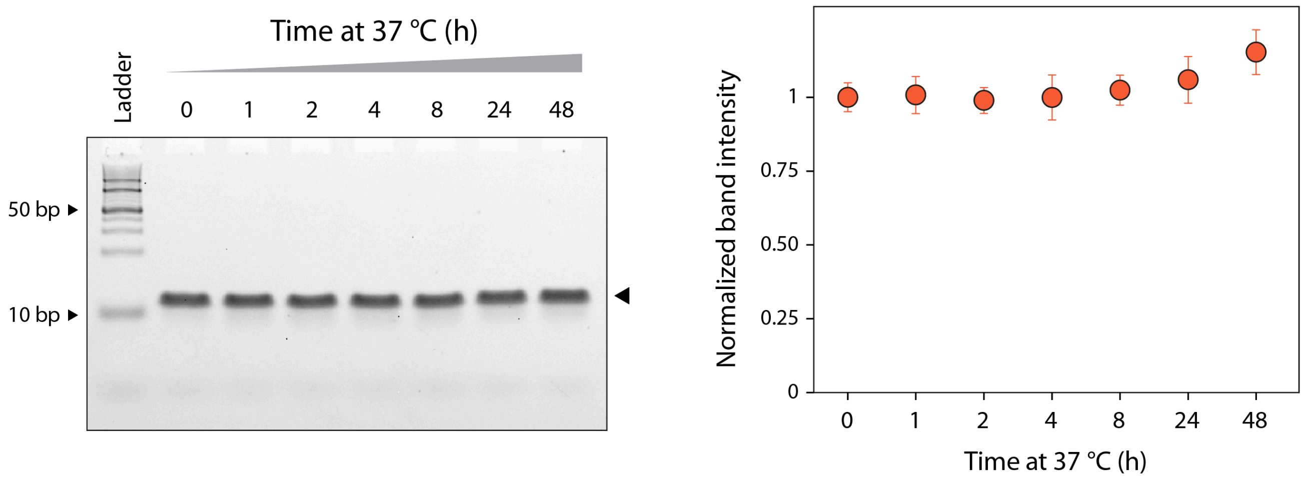


**Supplementary Fig. 20**. Stability of switchback DNA at 37 °C for up to 48 hours.

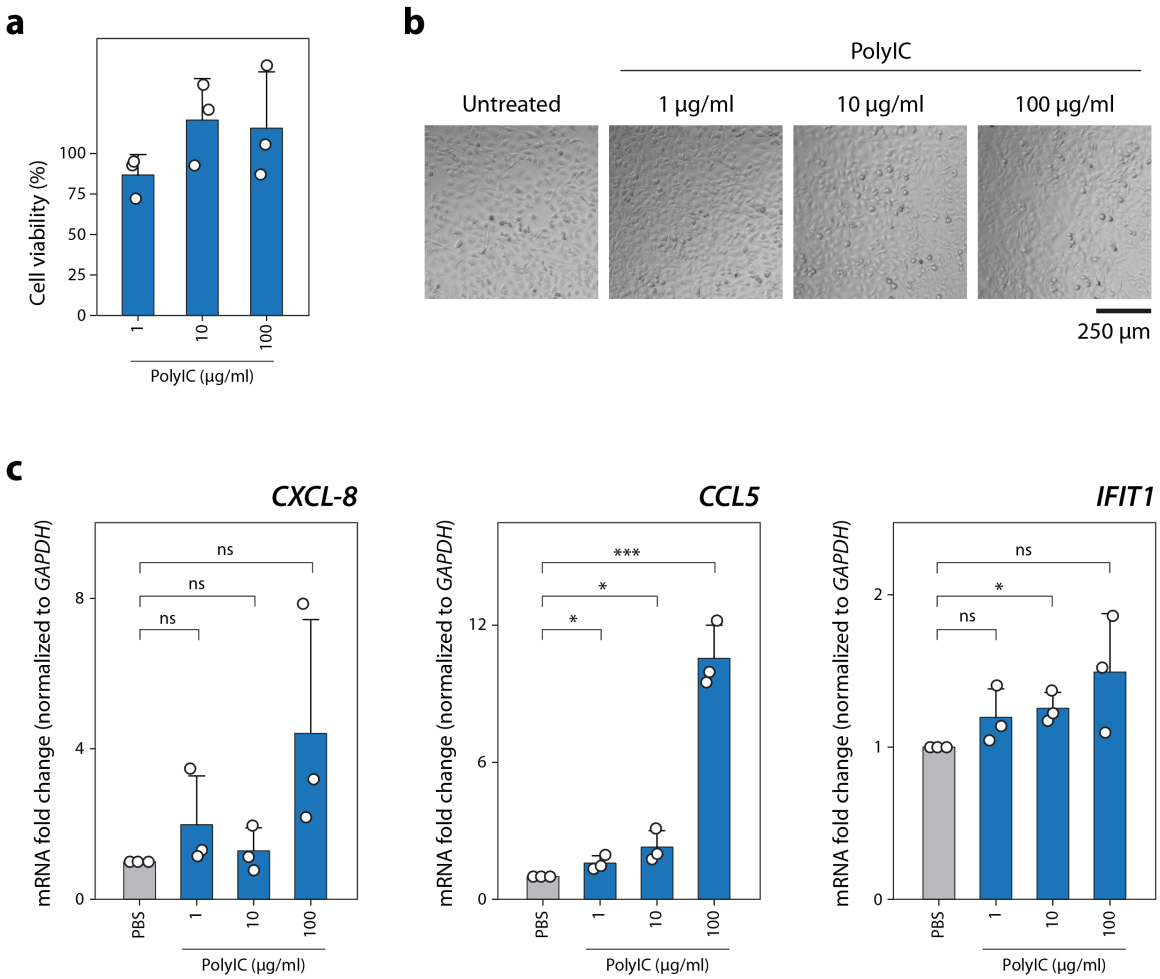


**Supplementary Fig. 21**. (a) Cell viability in HeLa cells incubated with different concentrations of polyIC. (b) Microscopy images of untreated cells and cells treated with polyIC. (c) RT-qPCR analysis of immune response markers. Data represent mean and standard deviation from biological triplicate experiments. Unpaired two-tailed *t*-test was used to compare polyIC treatment to PBS control, ns – not significant, $*P < 0.05$, $***P < 0.001$.

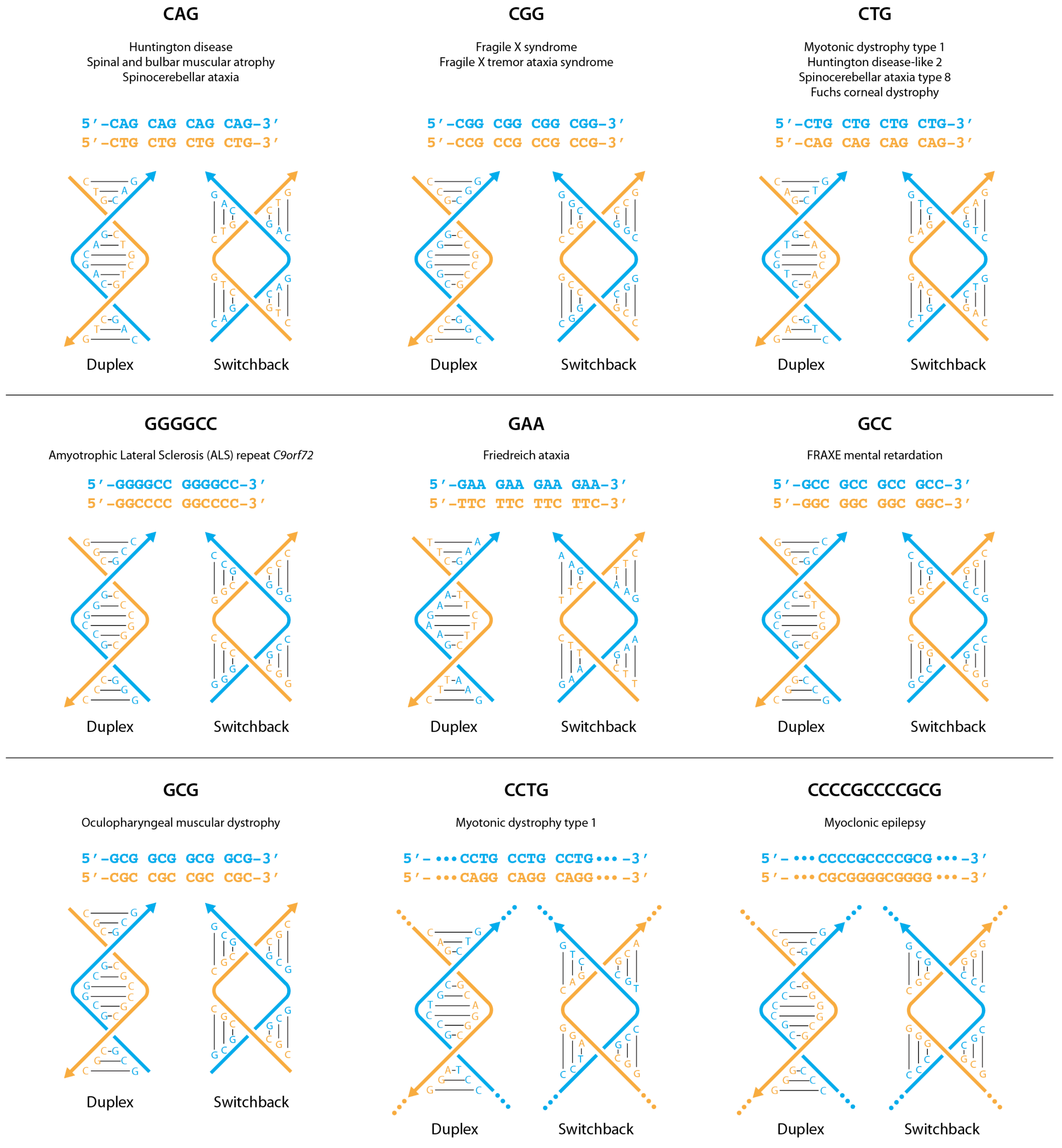


**Supplementary Fig. 22**. Select repeat sequences with the same set of complementary strands that are predicted to form either switchback DNA or a conventional duplex.

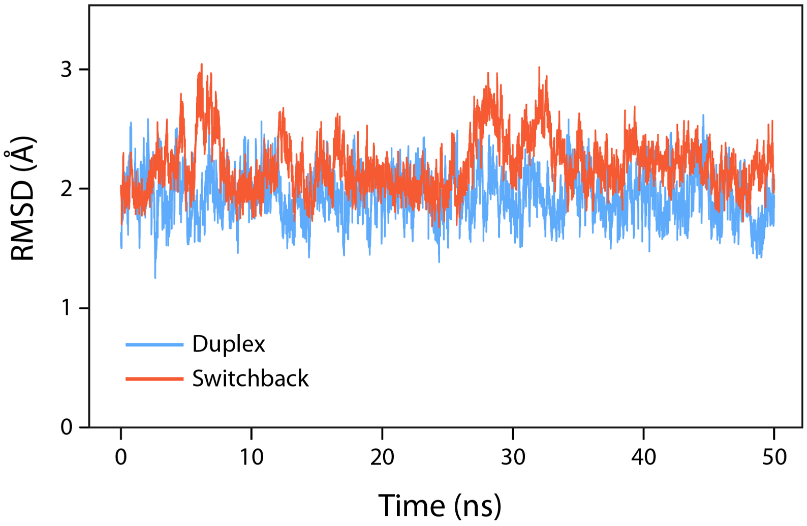


**Supplementary Fig. 23**. Root mean square deviation (RMSD) of the conventional duplex and switchback DNA as a function of time.

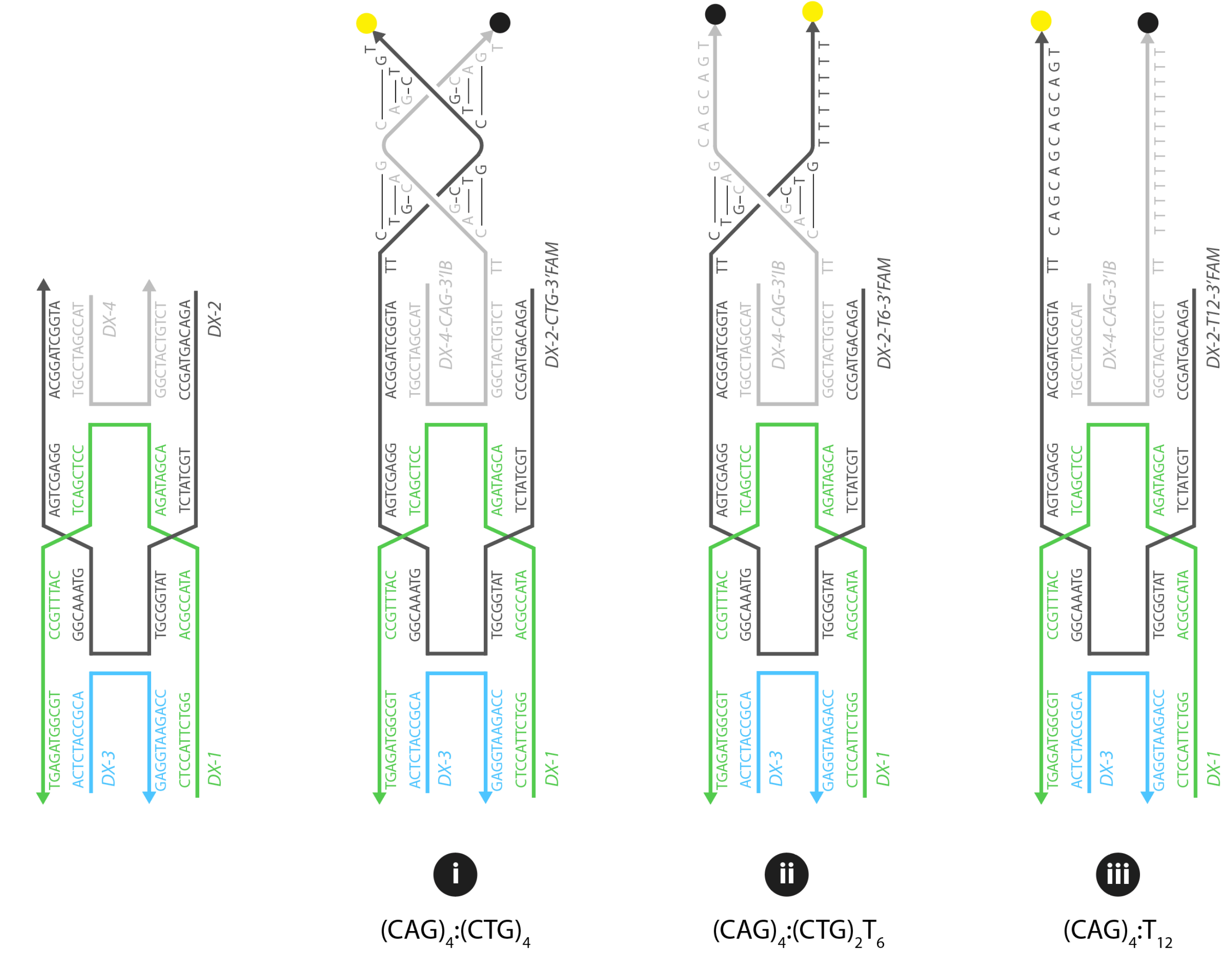

**Supplementary Fig. 24**. Design and sequences used in the DX scaffold.

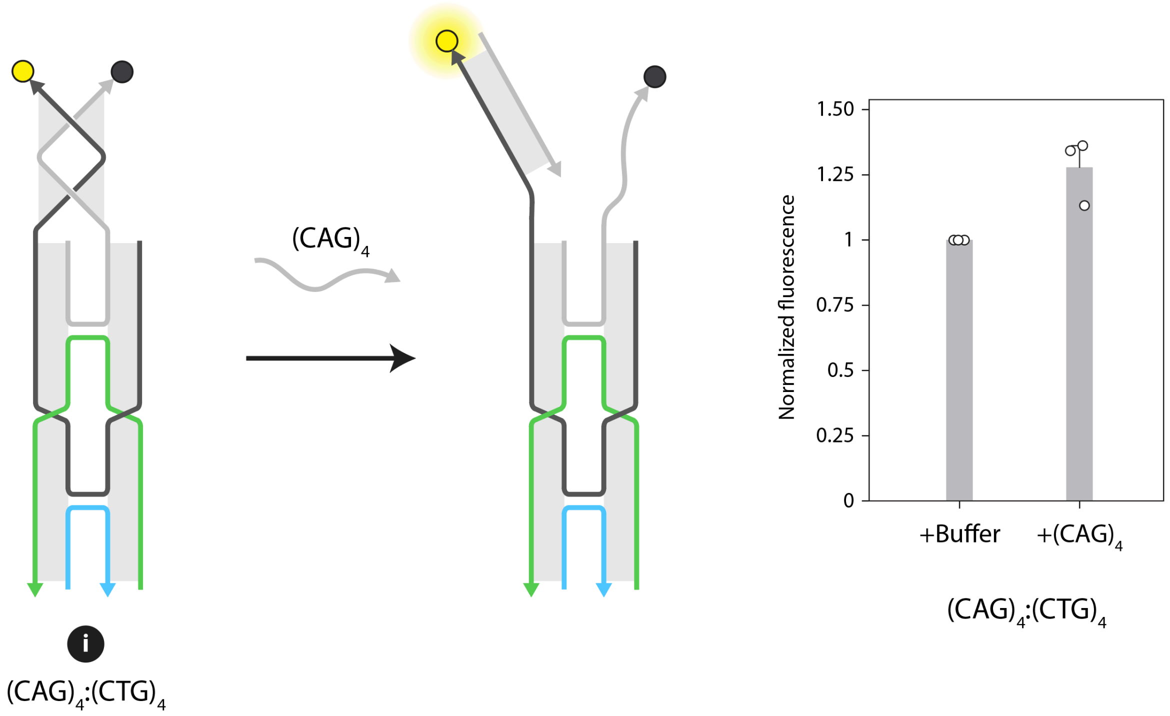


**Supplementary Fig. 25**. Fluorescence analysis of the interaction between two regions on the DX scaffold on the addition of a duplex complement.

**Supplementary Table 1.** Sequences used in this study. Strand combinations for different structures are shown in **Supplementary Fig. 1**, **9 and 24**.

| Name | Sequence (5′-3′) | Length |
|---|---|---|
| ***Homodimer switchback and corresponding duplex*** | | |
| A | TACGCGTGGATCCT | 14 |
| B | TGGATCCACGCGTT | 14 |
| B-long | TTTGGATCCACGCGTTTT | 18 |
| ***Heterodimer switchback and corresponding duplex*** | | |
| X | TACCAGCCGAACCT | 14 |
| Y | TGCTGGTGGTTCGT | 14 |
| Z | TGGTTCGGCTGGTT | 14 |
| Z-long | TTTGGTTCGGCTGGTTTT | 18 |
| X-5′FAM | /56-FAM/TACCAGCCGAACCT | 14 |
| Y-5′IB | /5IABkFQ/TGCTGGTGGTTCGT | 14 |
| Z-5′IB | /5IABkFQ/TGGTTCGGCTGGTT | 14 |
| ***Heterodimer switchback and corresponding duplex with mismatches*** | | |
| X-1m | TACGAGCCGAACCT | 14 |
| X-2mm-adj | TACGTGCCGAACCT | 14 |
| X-2mm-sep | TACGAGCCGATCCT | 14 |
| ***Control single strands and duplexes*** | | |
| C | TCTCAGTAAGCGTT | 14 |
| D | TACGCTTACTGAGT | 14 |
| C-long | TTTCTCAGTAAGCGTTTT | 18 |
| D-long comp | TTTACGCTTACTGAGTTT | 18 |
| ***Double crossover scaffold*** | | |
| DX-1 | CTCCATTCTGGACGCCATAAGATAGCACCTCGACTCATTTGCCTGCGGTAGAGT | 54 |
| DX-2 | AGACAGTAGCCTGCTATCTTATGGCGTGGCAAATGAGTCGAGGACGGATCGGTA | 54 |
| DX-3 | ACTCTACCGCACCAGAATGGAG TACCGATCCGTGGCTACTGTCT | 22 |
| DX-4 | TACCGATCCGTGGCTACTGTCT | 22 |
| DX-2-CTG-3′FAM | AGACAGTAGCCTGCTATCTTATGGCGTGGCAAATGAGTCGAGGACGGATCGGTATTCT GCTGCTGCTGT/36-FAM/ | 54 |
| DX-2-T6-3′FAM | AGACAGTAGCCTGCTATCTTATGGCGTGGCAAATGAGTCGAGGACGGATCGGTATTCT GCTGTTTTTTT/36-FAM/ | 54 |
| DX-2-T12-3′FAM | AGACAGTAGCCTGCTATCTTATGGCGTGGCAAATGAGTCGAGGACGGATCGGTATTTT TTTTTTTTTTT/36-FAM/ | 54 |
| DX-4-CAG-3′IB | TACCGATCCGTGGCTACTGTCTTTCAGCAGCAGCAGT/3IABkFQ/ | 22 |
| $(CAG)_4$ | TCAGCAGCAGCAGT | 14 |

**Supplementary Table 2.** ITC thermodynamic parameters of switchback and conventional duplexes.

| Parameter | Conventional duplex | Switchback DNA |
|---|---|---|
| Kd | 1.29 ± 0.8 nM | 30.73 ± 6.2 nM |
| n | 1.01 | 0.94 |
| $\Delta H$ | -83.33 ± 3.8 kcal/mol | -67.61 ± 1.8 kcal/mol |
| $-T\Delta S$ | 71.10 ± 4.2 kcal/mol | 57.35 ± 1.9 kcal/mol |
| $\Delta G$ | -12.23 ± 0.5 kcal/mol | -10.26 ± 0.1 kcal/mol |

**Supplementary Table 3.** Thermal melting analysis of structures containing mismatches.

| | Conventional duplex | | Switchback DNA | |
|---|---|---|---|---|
| | $T_m$ (°C) | $\Delta T_m$ (°C) | $T_m$ (°C) | $\Delta T_m$ (°C) |
| No mismatch | 66.5 ± 1.3 | - | 41.7 ± 1.0 | - |
| 1 mismatch | 58.5 ± 1.5 | 8.5 | 36.7 ± 1.5 | 5.0 |
| 2 mismatches (adjacent) | 54.3 ± 2.0 | 12.1 | 25.2 ± 0.3 | 16.5 |
| 2 mismatches (separated) | 44.3 ± 1.2 | 22.1 | 32.8 ± 0.8 | 8.8 |

**Supplementary Table 4.** Enzymes and reaction components used in biostability analysis.

| Nuclease | Cat # | Reaction components | Vendor |
|---|---|---|---|
| DNase I | M0303S | DNase I reaction buffer | New England Biolabs |
| T5 Exo | M0663S | NEBuffer 4 | New England Biolabs |
| Exonuclease V (Rec BCD) | M0345S | NEBuffer 4, ATP | New England Biolabs |
| Exonuclease VIII | M0545S | NEBuffer 4 | New England Biolabs |
| Micrococcal nuclease | M0247S | Micrococcal nuclease reaction buffer, recombinant albumin | New England Biolabs |

**Supplementary Table 5.** Calculated energy components that contribute towards the enthalpy change ($\Delta H$) calculation using the MMPBSA method, for the duplex and switchback DNA formed by CAG repeat sequences.

| Energy Component | Conventional duplex | Switchback DNA |
|---|---|---|
| $\Delta E_{VDW}$ | -61.07 ± 5.7 | -58.91 ± 6.1 |
| $\Delta E_{EL}$ | 2037.78 ± 38.4 | 2168.46 ± 66.9 |
| $\Delta E_{MM}$ | 1976.72 ± 37.6 | 2109.55 ± 65.4 |
| $\Delta G_{PB}$ | -2071.19 ± 36.3 | -2191.53 ± 62.5 |
| $\Delta G_{NP}$ | -6.41 ± 0.2 | -7.06 ± 0.3 |
| $\Delta G_{SOL}$ | -2086.5967 ± 36.4 | -2198.59 ± 62.6 |
| $\Delta H$ | -100.89 ± 4.8 | -89.04 ± 5.7 |

**Supplementary Table 6.** Primer sequences used for RT-qPCR.

| Gene | Forward | Reverse |
|---|---|---|
| *CXCL-8* | TGTCTGGACCCCAAGGAA | CATCTTCACTGATTCTTGGATACC |
| *CCL5* | GAGTATTTCTACACCAGTGGCA | GACTCTCCATCCTAGCTCATCT |
| *IFIT1* | CCACAAGACAGAATAGCCAGAT | GCTCCAGACTATCCTTGACCT |
| *GAPDH* | CACGTTTTGGATGCACTGAGAC | GATGGAGGGCCTTTTATTCGCG |
